# Introducing an *rbc*L and *trn*L reference library to aid in the metabarcoding analysis of foraged plants from semi-arid eastern South African savannas

**DOI:** 10.1101/2022.10.06.511093

**Authors:** Danielle Botha, Mornè du Plessis, Frances Siebert, Sandra Barnard

## Abstract

The success of a metabarcoding study is determined by the extent of taxonomic coverage and the quality of records available in the DNA barcode reference database used. This study aimed to create an *rbc*La and *trn*L (UAA) DNA barcode sequence reference database of plant species that are potential herbivore foraging targets and commonly found in semi-arid savannas of eastern South Africa. A study-area-specific species list of 755 species was compiled. Thereafter, reference libraries for *rbc*La and *trn*L (UAA) sequences were created mined from sequence databases according to specific quality criteria to ensure accurate taxonomic coverage and resolution. The taxonomic reliability of these reference libraries was evaluated by testing for the presence of a barcode gap, identifying a data-appropriate identification threshold, and determining the identification accuracy of reference sequences via primary distance-based criteria. The final *rbc*La reference dataset consisted of 1238 sequences representing 318 genera and 562 species. The final *trn*L dataset consisted of 921 sequences representing 270 genera and 461 species. Barcode gaps were found for 76% of the taxa in the *rbc*L barcode reference dataset and 68% of the taxa in the *trn*L barcode reference dataset. The identification success rate, calculated with the *k*-nn criterion was 85.86% for the *rbc*L dataset and 73.72% for the *trn*L dataset. The datasets for *rbc*L and *trn*L combined during this study are not presented as a complete DNA reference library, but rather as two datasets that should be used in unison to identify plants present in the semi-arid eastern savannas of South Africa.

## 1. Introduction

Dietary analysis is a fundamental part of constructing habitat selection and utilization models as well as assessing the influence of land use type on the plant community and how this, in turn, can influence foraging strategies [1, 2]. Determining and analysing food items in ecosystems will also aid in identifying key environmental resources for the design of reliable conservation and management strategies [3, 4]. To determine the composition of animal diets and how this composition reflects the plant community in question, animal faeces can be examined to identify food items [2,5], as well as the relative abundance thereof ingested per individual sampled [6]

The metabarcoding approach allows for species identification from heterogeneous and degraded environmental samples based on the amplification and subsequent analysis of DNA barcodes [7,8], which are short gene sequences from a standardized region of the genome that displays low intraspecies and high interspecies variability [9,10]. DNA metabarcoding provides an attractive alternative to the traditional methods of large-scale species identifications and it can be applied to microscopic [11], cryptic [12], and digested materials [2,13].

The r*bc*La and *trn*L (UAA) DNA barcodes have been proposed as two single-locus gene regions that can be used for the analysis of herbivorous- and omnivorous animal diets via DNA metabarcoding [2,13–16]. These two barcodes were chosen as the core barcoding regions for this study due to their coverage of different savanna plant species and because of their sequence availability on global databases.

Except for the choice of the barcoding marker to be used, the efficiency and usefulness of metabarcoding as a tool are underpinned by the extent of taxonomic coverage and the quality of records available in the DNA barcode reference database to which the query sequences will be compared and identified [17,18]. However, there is a lack of plant taxonomic sequence information for some ecological regions on global DNA sequence databases [19]. According to [19], as of 2019 DNA sequences available on the Barcode of Life Data (BOLD) database represent only ~8% of the vascular plants in Africa, while Africa compromises ~20% of the earth’s landmass and owns approximately 15% of the global plant diversity [20]. Furthermore, forbs (i.e., non-graminoid herbaceous vascular plants: [21,22], which contribute more than 70% to herbaceous species richness in grasslands, temperate deciduous forests, and savanna ecosystems, have been neglected in vegetation studies and knowledge of their ecological role in savanna ecosystems are limited [23]. As a result, forbs are also weakly represented on global DNA sequence databases.

Furthermore, it is well known that public DNA reference databases may contain sequencing errors and unverified taxonomies by a lack of voucher specimen information and links to metadata. This is especially true for GenBank where the curation of the database is left largely to scientists submitting sequences [24,25]. Since the DNA reference database will determine the accuracy of taxonomic assignations i.e., the success of a barcode in distinguishing between species [26], it is necessary to create a subset of the data available on public DNA databases that are manually curated for sequence quality, taxonomic validity, geographic validity as well as to test the efficiency of this subset reference database to discriminate between and within species with similarity, distance- and tree-based methods.

With this study we therefore aim to: (i) create a comprehensive *rbc*La and *trn*L (UAA) DNA barcode reference database of plant species commonly found in a semi-arid South African savanna, and (ii) evaluate the efficiency of this reference database in discriminating between and within species by testing for the presence of a barcode gap, identifying a data appropriate identification threshold as well as determining the identification accuracy of reference sequences via primary distance-based criteria. The fulfilment of these aims will result in a DNA reference database that can be used confidently to delineate between species found in the faeces of herbivores foraging in a semi-arid eastern South African savanna as well as to use the successful taxonomic identifications to make certain deductions about the plant community to aid in management and conservations strategies.

## 2. Materials and methods

### 2.1. Barcode reference list assembly

The compilation of this DNA reference database of species found in a semi-arid eastern South African savanna is challenging since scientific publications listing all taxa likely to be encountered in the diets of herbivores foraging in these ecosystems were largely unavailable. The semi-arid eastern savannas of South African, located in the Limpopo and Mpumalanga provinces is defined as savannas receiving a mean annual precipitation of below ~650 mm rendering them ‘unstable’, therefore needing disturbances such as fire and herbivory to maintain the co-existence of trees and grasses [27]. The species lists used as reference for sequence mining was built around two sites at which floristic surveys have been undertaken for the purpose of community assemblage changes with grazing [28]. Dominant plant species recorded at these sites were sequenced by the North-West University (NWU) if they were poorly represented in DNA sequence libraries. Syferkuil is an experimental farm in the Limpopo province, and Welverdiend (Mpumalanga province) a rural settlement with open spaces for communal grazing. To expand the reference database accuracy this list was also enriched with applicable floristic and DNA sequencing data from other studies based in African savannas from which DNA sequences were available (e.g. [2,19]. A subset of *rbc*La and *trn*L (UAA) records available on public databases was created by using the compiled species list (Supplementary Table 1) to mine sequences according to specific quality criteria to ensure accurate taxonomic coverage and resolution. These criteria included sourcing a maximum of three individuals per species per marker, not including sequences from the same herbarium, and giving preference to sequences with voucher specimens and sequences with the least ambiguous bases. Accession numbers of barcodes that match the quality criteria and have been sequenced with the *rbc*L primers, *rbc*La-F: ATGTCACCACAAACAGAGACTAAAGC [29] and *rbc*La-R: GTAAAATCAAGTCCACCRCG [30] as well as with the *trn*L primers [31], *trn*L-c: CGAAATCGGTAGACGCTACG and *trn*L-d: GGGGATAGAGGGACTTGAAC, can be found in Supplementary Table 2. The public databases used to source sequences in this study were the GenBank nucleotide database (www.ncbi.nlm.nih.gov) and the BOLD System public database (www.boldsystems.org).

**Table 1:**
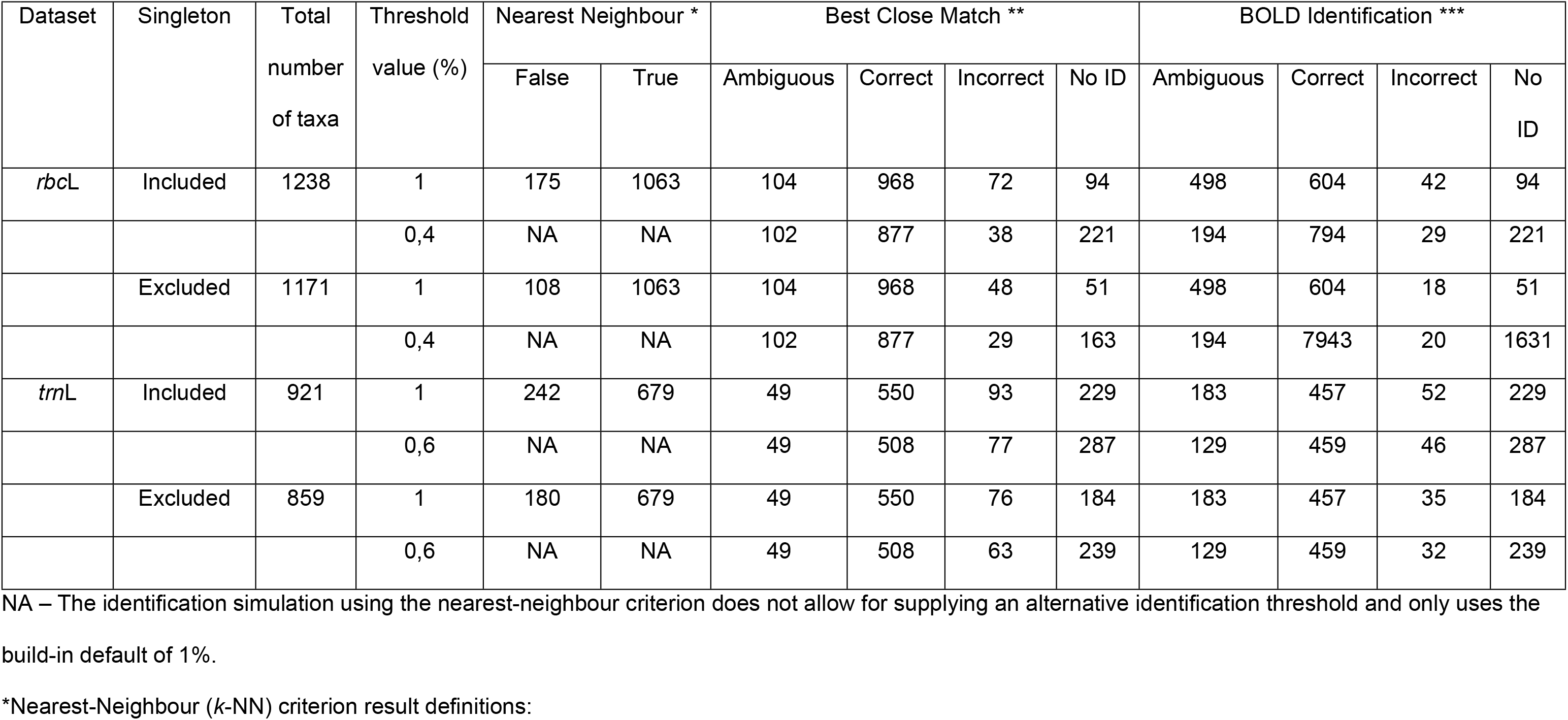
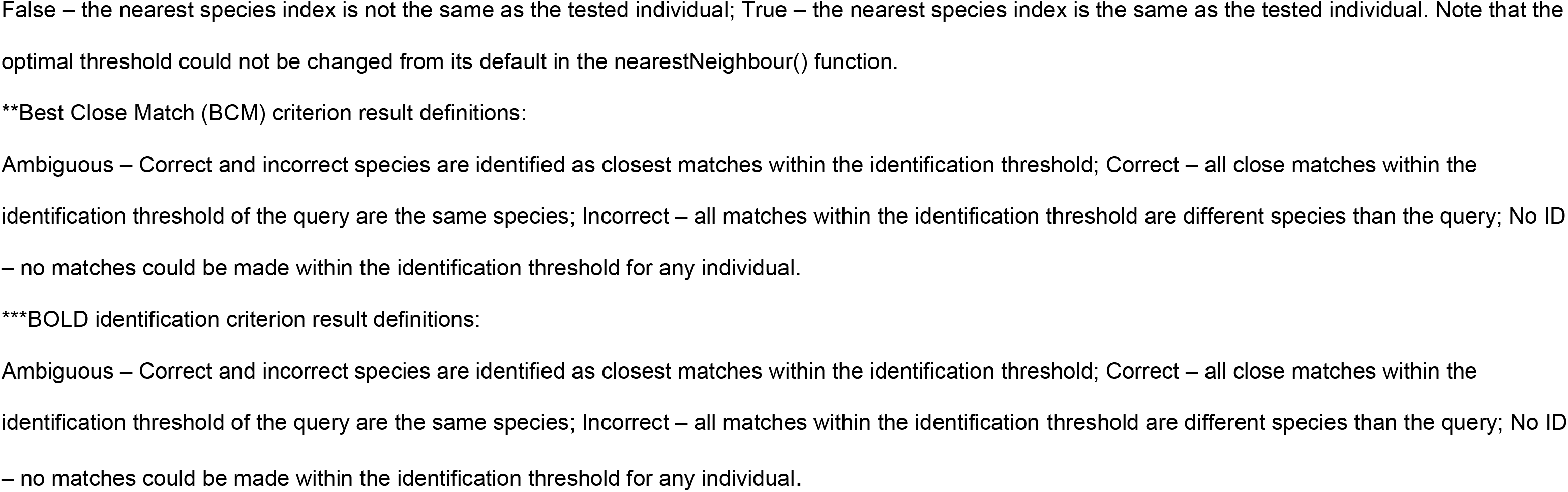
Discriminatory power of the *rbc*L and *trn*L reference datasets predicted by the application of three distance-based measures: nearest neighbour, Meier’s best close match, as well as BOLD identification criterion, for both default and optimized thresholds for the inclusion and exclusion of singletons for the respective datasets. All three analyses were performed to identify queries within the respective identification thresholds to genus level.

| Dataset | Singleton | Total<br>number<br>of taxa | Threshold<br>value (%) | Nearest Neighbour * |  | Best Close Match ** |  |  |  | BOLD Identification *** |  |  |  |
| --- | --- | --- | --- | --- | --- | --- | --- | --- | --- | --- | --- | --- | --- |
|  |  |  |  | False | True | Ambiguous | Correct | Incorrect | No ID | Ambiguous | Correct | Incorrect | No<br>ID |
| *rbcL* | Included | 1238 | 1 | 175 | 1063 | 104 | 968 | 72 | 94 | 498 | 604 | 42 | 94 |
|  |  |  | 0,4 | NA | NA | 102 | 877 | 38 | 221 | 194 | 794 | 29 | 221 |
|  | Excluded | 1171 | 1 | 108 | 1063 | 104 | 968 | 48 | 51 | 498 | 604 | 18 | 51 |
|  |  |  | 0,4 | NA | NA | 102 | 877 | 29 | 163 | 194 | 7943 | 20 | 1631 |
| *trnL* | Included | 921 | 1 | 242 | 679 | 49 | 550 | 93 | 229 | 183 | 457 | 52 | 229 |
|  |  |  | 0,6 | NA | NA | 49 | 508 | 77 | 287 | 129 | 459 | 46 | 287 |
|  | Excluded | 859 | 1 | 180 | 679 | 49 | 550 | 76 | 184 | 183 | 457 | 35 | 184 |
|  |  |  | 0,6 | NA | NA | 49 | 508 | 63 | 239 | 129 | 459 | 32 | 239 |

NA – The identification simulation using the nearest-neighbour criterion does not allow for supplying an alternative identification threshold and only uses the build-in default of 1%.
\*Nearest-Neighbour (*k*-NN) criterion result definitions:
False – the nearest species index is not the same as the tested individual; True – the nearest species index is the same as the tested individual. Note that the optimal threshold could not be changed from its default in the nearestNeighbour() function.
\*\*Best Close Match (BCM) criterion result definitions:
Ambiguous – Correct and incorrect species are identified as closest matches within the identification threshold; Correct – all close matches within the identification threshold of the query are the same species; Incorrect – all matches within the identification threshold are different species than the query; No ID – no matches could be made within the identification threshold for any individual.
\*\*\*BOLD identification criterion result definitions:

This study also enriched the sourced dataset by supplying twenty-four sequences from plants found in semi-arid eastern South African Savanna ecosystems. Most of these sequences are known to not occur at all, or not frequently on public DNA sequence databases. A pipeline for the collection, DNA isolation, and sequencing of *rbc*La and *trn*L barcodes has been established in 2015. Leaf material and voucher specimens of woody and herbaceous plants were collected from Welveridend, Syferkuil and neighbouring landscapes. Botanists at the NWU examined the voucher specimens to confirm their taxonomy and they were entered into the local Goossens Herbarium of the NWU. The barcode sequences were added to the GenBank nucleotide database and included in the reference database compiled and evaluated in this study. GenBank accessions numbers for the *rbc*La barcode range from MZ461574.1 to MZ461594.1 and the *trn*L barcode from MZ461547.1 to MZ461570.1. (Supplementary Table 3)

### 2.2. Phylogeny

The candidate barcode reference sequences for both markers to be used for phylogenetic analysis were respectively aligned using Geneious (v. 2021.2.2) via MUltiple Sequence Comparison by Log-Expectation (MUSCLE) [32], with a maximum number of eight iterations. The sequences for the *trn*L marker were not split into families, as they will be for downstream analysis, due to the requirement of robust representations of the alignment and clusters formed for the subsequent “quality” analysis.

The aligned candidate barcode reference sequences were used to reconstruct neighbour-joining (NJ) phylogenetic trees for the *rbc*La and *trn*L dataset, as implemented by Geneious with bootstrap testing of 1000 replicates using the Jukes-Cantor model. Within angiosperms, *Amborella* is reported as the sister to all remaining flowering plants [33,34], and accordingly, the *rbc*La and *trn*L (UAA) genes of *Amborella trichopoda* were included as an outgroup for the *rbc*La and *trn*L sequences and aligned with the rest of the datasets. The individuals in both the *rbc*La and *trn*L phylogenetic tree were also reviewed manually to validate clusters and sub-clusters that were formed. This evaluation was put in place to improve further analyses recommending the removal of sequences showing an evident deviation from their cluster, be it a cluster of life forms (such as trees, shrubs, grasses, forbs) or plant families. The sequences that were identified to have problematic placements were then replaced with another individual sequence of the species with equal quality if possible, or it was removed from further analysis.

### 2.3. DNA barcode assembly for reference database

DNA sequences for both marker datasets were exported to RStudio and aligned to prepare sequences for downstream analysis. Alignment was again facilitated with MUSCLE, but without the outgroup sequences, and subsequent determination of genetic distance and barcode-gap analysis took place. All sequences for *rbc*La were aligned simultaneously using the *AlignSeqs()* function in the DECIPHER package [35], and *trn*L sequences were split into families [19] and accordingly aligned with the same function due to the often highly divergent sequences displayed for distantly related species. Distance matrices for both datasets were calculated using the Kimura-2 parameter (K2P) model [36] which is the build-in model of evolution in the APE package’s *dist.dna()* function [37]. Paradis & Schliep

### 2.4. Barcode gap analysis

The barcode analysis for each marker dataset used the K2P model to calculate genetic distances for each sequence per marker and was done using the R package SPIDER (v.1.5) [38]. A barcode gap analysis is performed by two approaches, as recommended by [39], which will allow for barcode gap analyses of the complete datasets, as well as barcode gaps for individual sequences represented in the respective datasets. The first approach compares the median of inter – and intraspecific distances for each marker dataset and assesses the significance of the differences between genetic distances with a Wilcoxon rank-sum test with the R function *wilcox.test()* in the R STATS package. The second approach follows the determination of the presence of a barcode gap following [40], where the difference between maximum intraspecific distance and the minimum interspecific distance is calculated for each sequence per marker dataset. Both approaches were applied during this study.

### 2.5. Analysis with distance-based criteria

The optimisation of the species identification threshold for each marker dataset is performed to increase identification success [26]. During this study, we used the *threshOpt()* function in the SPIDER package of R (v.1.5) [38]: to systematically subject the marker dataset to a range of threshold values (between 0.1 to 0.2% K2P distance). The optimal identification threshold was then calculated based on the number of true positive, false positive, true negative, and false negative identifications for each threshold in the range. The optimal threshold was identified as the value predicting the lowest cumulative error, i.e., the lowest amount of false-positive and negative identifications according to [38].

The predicted accuracy of taxonomic assignation of the reference sequences was determined using three distance-based analyses via the SPIDER package of R (v.1.5) [38]. These primary distance-based criteria included the nearest neighbour (*k*-nn) criterion [41], Meier’s best close match (BCM) criterion [42] as well as BOLD identification criterion [43]. The functions by which these measures are executed (*nearNeighbor()*, *bestCloseMatch()*,and *threshID()*, respectively performed an internal taxonomic assignation, where individual sequences were treated as unknown queries and the rest of the marker dataset was treated as the DNA reference database to match corresponding taxonomies within the identified optimal threshold.

## 3. Results

### 3.1. DNA reference dataset: phylogenetic analysis and summary statistics

The final species list (Supplementary Table 1) consists of 755 species, 356 genera, 99 families as well as 3 classes (Magnoliopsida, Liliopsida, and Polypodiopsida) in the phylum Tracheophyta. Of these 755 species, sequences for 108 could not be found for either the *rbc*L or *trn*L barcode. Upon inspection of the NJ trees of rbcL and *trn*L, it was advised to replace 265 individuals with other sequences from the same *species* of equal quality since they did not group with other members of their family or life form. Sequences for 59 individuals could be replaced, but 206 sequences had to be removed. This led to a loss of taxonomic coverage in some species, now only represented by < 3 individuals and the complete loss of 57 unique species from the reference dataset. The remaining species and accession numbers, or process identities (process IDs) in the case of BOLD sequences, can be found in the supplementary table which now constitutes the complete DNA reference database. The composition of the complete reference database is illustrated in Figure 1, where the family Poaceae is the largest representative with 47 genera and 109 species. The second-largest representative family is the Fabaceae with 35 genera and 87 species. The *rbc*La reference dataset consists of 1238 sequences representing 318 genera and 562 species. The *trn*L dataset consists of 921 sequences which represent 270 genera and 461 species. Naturally, there is an overlap between the datasets as sequences were mined for the same species for both markers: sequences for 345 species of 224 genera are found in both marker datasets.

**Figure 1.**
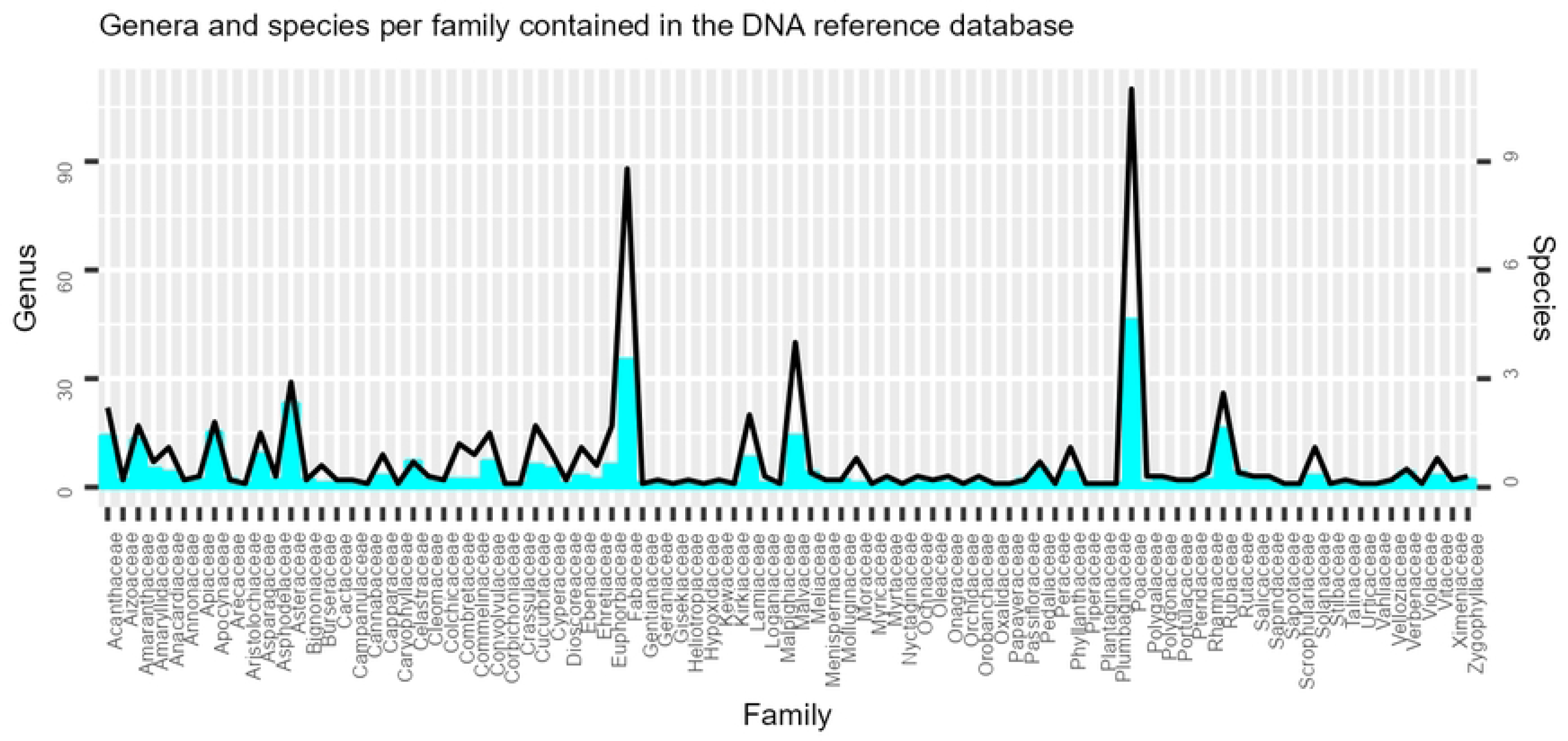
The composition of the final DNA sequence reference database in terms of number of genera (histogram) and species (line plot) per family.

### 3.2. Reference dataset analysis

According to the first approach of the barcode gap analysis, by which the presence of a barcode was assessed for both marker datasets by the comparisons of medians for inter-and intraspecific genetic distances, it was found that for the *rbc*L dataset, the interspecific K2P distances ranged from 0,00 – 0,94 with a median of 0.01. The interspecific distances were statistically larger than the intraspecific K2P distances (Wilcoxon test: P < 2.2e −16) with ranges of 0 – 1.45 and a median of 0.00. The *trn*L dataset followed the same trend with generally larger interspecific distances (ranges from 0,00 – 1,01 and a median of 0,01) than intraspecific distances (ranges: 0 – 1,07; median: 0; Wilcoxon test: p < 2.2e −16). The second approach [42], revealed a positive difference between maximum intra- and minimum interspecific K2P distances for 76% of the *rbc*L sequences and 68% for the *trn*L sequences. These results are reported as scatterplots (Fig. 2) where the sequences that showed a gap appear above the 1:1 slope.

**Figure 2:**
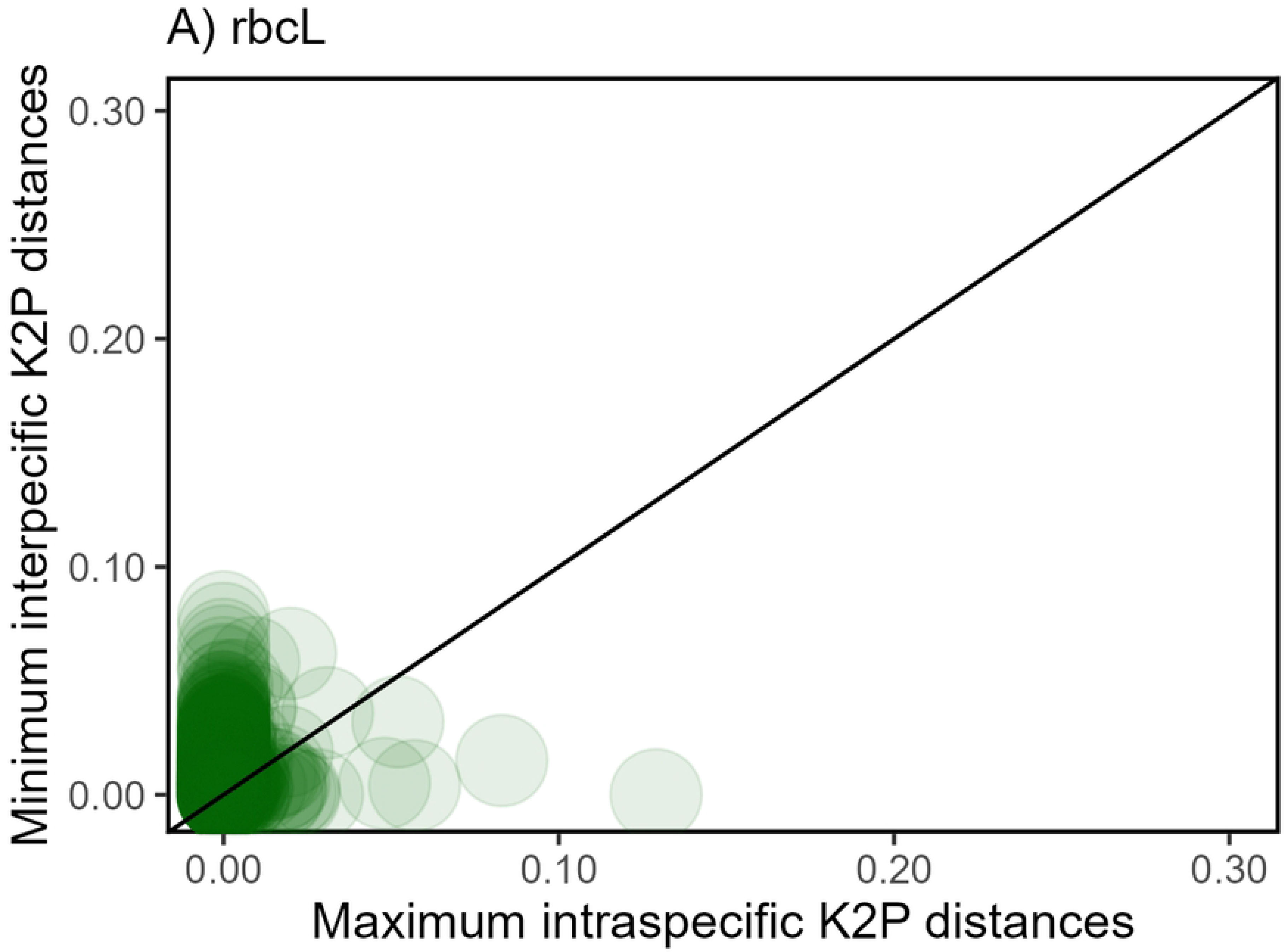
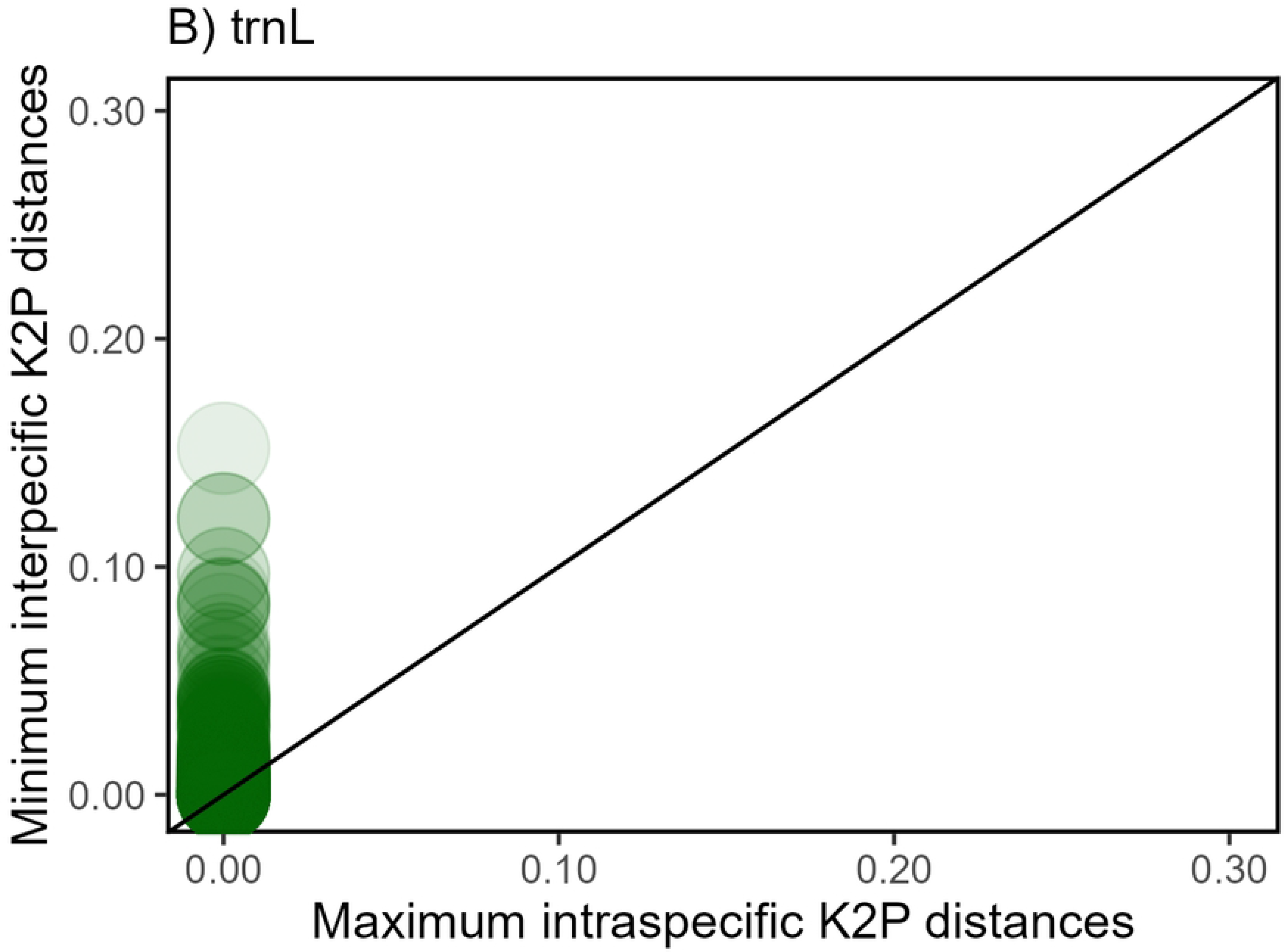
Scatterplots of the Barcode gap s for species present A) in the *rbc*L dataset and (B) *trn*L dataset. Scatterplots show the relationships between minimum interspecific distance (y-axis) and maximum intraspecific distance (x-axis) where the barcode gap is indicated by points falling above the 1:1 slope. High densities of plots are indicated by the darker colours.

To improve the accuracy of the DNA reference database, optimal identification thresholds were determined with the threshOpt function in the R package SPIDER. The default threshold of species optimisation used by identification algorithms such as BLAST and BOLD is 1%, which may not always be appropriate for all reference datasets. Accordingly, the determined optimal threshold for the *rbc*L dataset was 0.4% and that of the *trn*L dataset was 0.6%, which clearly indicates that the identification accuracy of the datasets would not have been accurately portrayed by the default threshold value of 1%. These optimal threshold values will be used during the simulated taxonomic identification via the three distance-based analyses which will aid in predicting the accuracy of the reference datasets.

The nearest-neighbour (*k*-NN) criterion performed the best among the distance-based measures during the identification simulations (Table 1). The identification success rate, calculated as the number of matches that share the same species index as the query was 85.86% for the complete *rbc*L dataset and 73.72% for the *trn*L dataset. These identification success rates can be increased by excluding singletons, resulting in an identification success rate of 90.78% and therefore an increase of 4.92% for the *rbc*L dataset. Excluding singletons led to an increase of 5.33% resulting in a success rate of 79.05% for the *trn*L dataset. Meier’s [41] best close match (BCM) criterion performed similarly to the *k*-NN identification criterion with an identification success rate of 82.66% correct matches with the optimized threshold value for the exclusion of singletons for the *rbc*L dataset and 64.03% for the *trn*L dataset with matches performed in the optimized threshold with the exclusion of singletons. The BOLD identification criterion failed to show the same level of identification success rates with the inclusion of an optimized identification threshold. The BOLD identification criterion performed the worst among the other measures with the highest discriminating power shown for the *trn*L dataset with the exclusion of singletons, reaching 53.20% with the default threshold value of 1%.

## 4. Discussion

### Reference library development

Despite several studies in which DNA barcoding was applied in the analyses of herbivore diets [3,44,45], a focus on herbivore diets in African savannas remains limited [2, 19]. Until present, none focused on South African savannas, and there is not currently a robust DNA reference sequence library available for the semi-arid eastern South African Savanna. During this study we have curated a DNA reference library consisting of both and the *rbc*L and a *trn*L dataset. This is the first step towards facilitating metabarcoding studies aimed at South African savanna ecosystems, specifically for the identification of plant materials obtained in herbivorous faeces.

The ideal construction of the reference library would be to sample, identify, and sequence the relevant barcodes of all the species on the species list. However, this would be a costly and laborious research campaign. Mining sequences from global databases is a much more cost-effective option and the method illustrated here can be reproduced for any nature of metabarcoding study aimed at representing the composition of an ecosystem without manually sequencing all specimens. The use of GenBank as a taxonomic resource has been questioned since there exists an absence of preserved voucher specimens, non-justified species identifications, and low-quality data [42,46]. The same is true for BOLD, although this database is better curated due to higher quality submission standards but may still contain incorrectly identified taxa and low-quality sequence data [42,43]. Being aware of these limitations enables the user to create certain download criteria which result in a database of sequences with known localities, known voucher specimens as well as the inclusion of barcodes with the least ambiguous bases. This database, as well as any database mined from global reference databases, is not impervious to imperfection, but this method of using a species list and applying download quality criteria will certainly lift its standard above a taxonomic assignation against whole databases such as GenBank and BOLD. The lowest level of taxonomic resolution achieved in metabarcoding studies is typically genus-level [44,47]; Turunen *et al*., 2021) and the assignation of taxonomy at lower taxonomic levels is often infeasible and inaccurate. Therefore, to ensure that this database of barcode sequences is robust enough to be used as an identification tool the barcode gap was analysed together with the barcode phylogeny.

Disparities between inter-and intraspecific distances between and among species are defined as a barcoding gap that, if present at a locus, enables the reliable differentiation of species [26, 38]. A barcode gap was evident in both approaches used and for both datasets of which the reference library consists of. A barcode gap for 76% of the species was obtained for the *rbc*L barcode sequences and for 68% of the species for the *trn*L sequences. This implies that reliable species differentiation would not be possible for24% of the species in the *rbc*L dataset and 32% in the *trn*L dataset. However, limitations have also been reported in other DNA-barcode studies such as [19, 48and 30]. Gill *et al*., (2019) [19] reported a barcode gap for only 73% of the species investigated for the *rbc*L primer and 79% for the *trn*L-F primer in their barcode library for the semi-arid eastern South African Savanna plants. Mishra *et al*., (2017) [48] have shown that even within one genus (*Terminalia*) the three different barcodes of *mat*K, ITS and *rbc*L contained a barcode gap for <70% of the species. As stated by Gill *et al*., (2019) [19] savannas are comprised of a diverse range of species that are prone to the absence of a barcoding gap.

The distance-based measures of *k*-nn, BCM, and BOLD identification criteria infer identification success rates by considering the K2P distance matrix of each dataset to simulate taxonomic identifications of one sequence against the rest of the databases to match taxonomies within the identification threshold [37]. Usually, the distance-based matrices are applied at a species level, but for this metabarcoding study, it was decided to predict the identification success of the datasets based on genus-level identifications (Table 1). The best success rates across all three methods were seen with the exclusion of singletons, in this case, genera represented by only one individual. The analysis of the identification of success rates, with the exclusion of singletons, is also a prerequisite for the alignment that precedes the barcode gap analysis. Singletons are typically a problem in barcoding studies since the identification simulations will treat the singleton as a query, and it will not have a match available in the reference dataset and will result in either “incorrect” or “no identification” [37].

Analyses revealed that the *rbc*L dataset has a predicted identification success rate of 90.78% and *trn*L 79.05% with the *k*-NN method. This would imply that a barcode gap analysis is not always an accurate predictor of the species discrimination success of sequences in a reference database, as concluded in certain studies [40,49, 50]. This is also reflected by the findings in this study as illustrated by the inconsistencies between identification success rate simulations and the evaluation of the barcode gap. However, in this case, the barcode gap analysis can be accepted as a more efficient tool to predict database accuracy since it displays the identification success rate achieved by species-level identifications, whereas the distance-based analysis implemented genus-level taxonomic assignments. Furthermore, the discrimination success rates demonstrated by both datasets are above (*rbc*L = 76%) or very close (*trn*L = 68%) to the discrimination success rate of 72% as proposed by the CBOL Plant Working Group (2009) [51]. This implies that an adequate discrimination success rate would be possible to the lowest taxonomic level of genus when taking the predicted identification success rates into account.

The phylogenetic analysis served as a re-identification strategy of sequences included in the *rbc*L and *trn*L reference database. The exclusion of singletons, as recommended by the identification rates of the various distance-based measures (Table 1) was not considered in this part of the study, as this would have led to a great loss of taxonomic coverage and identifications as seen by a drop of 5.41% and 7.22% in the total taxonomy included in the *rbc*L and *trn*L databases, respectively. Excluding singletons from the reference datasets would lead to a loss of the families of flowering plants, namely: *Aristolochiaceae*, *Dioscoreaceae*, *Gentianaceae*, *Kirkiaceae*, *Passifloraceae*, *Piperaceae*, *Plantaginaceae*, *Urticaceae* in the *rbc*L datasets as well as the loss of *Hypoxidaceae*, *Menispermaceae*, *Nyctaginaceae*, *Passifloraceae*, *Piperaceae*, *Rutaceae*, *Scrophulariaceae* and *Stilbaceae* in the *trn*L dataset. We, therefore, sacrificed some identification success for the inclusion of singletons, and therefore an accurate representation, in terms of species present, in a South African savanna. The NJ trees revealed that sequences formed cohesive clusters of orders for the *rbc*L dataset. However, an obvious low taxonomic resolution for some orders in the *trn*L dataset was again obvious, such as for the orders Poales and Commelinales and Fabales. Similar results were also shown by [19] for the families Poaceae, Fabaceae, and Malvaceae. Low taxonomic resolutions were expected since many genera which are known to occur in South African savannas are under-sampled in global reference databases or are yet to be barcoded with either *rbc*L or *trn*L, leading to a lack of representatives for certain genera or the inclusion of singletons in the reference database. Species represented in the South African savannas are very diverse as can be seen from the sum of the branch lengths of the NJ trees, namely 108.23 for the 922 *trn*L-barcode taxa and 382.34 for the 1239 *rbc*L-barcode taxa.

### Conclusion

During this study, we developed a DNA barcode reference library that is robust to identify taxons on the list of species that we curated for some foraged plants from the semi-arid eastern South African savanna. All the different tests used to validate the use and accuracy of this library indicate that it can be used with confidence to assign taxonomies for plants found in the eastern savannas of South Africa. We envisage that it will add to other similar research done not only on the local flora but also to the work done on savannas elsewhere on the African continent. The datasets for *rbc*L and *trn*L are not presented as a complete DNA reference library, but rather as two databases that should be used in unison to identify species foraged on in the semi-arid eastern South African Savanna.

## Acknowledgments

We thank Miss Siena Bieler for assistance with field collections, DNA isolation, and sequencing of barcodes. We thank the North-West University for its support of this research.

## Notes

### Competing Interest Statement

The authors have declared no competing interest.

